# Perisurgical Colony Stimulating Factor 1 (CSF1) treatment ameliorates liver ischaemia/reperfusion injury in rats

**DOI:** 10.1101/2025.03.12.642926

**Authors:** Sarah Schulze, Sahar Keshvari, Gregory C. Miller, Kim R. Bridle, David A. Hume, Katharine M. Irvine

## Abstract

**Background:** In the context of hepatobiliary and liver transplant surgery, ischemia-reperfusion (I/R) injury can occur due to temporary interruption of blood flow to the organ followed by a potentially damaging inflammatory response to reperfusion. As macrophages can promote liver growth and contribute to resolution of chronic liver injury and fibrosis, we tested the hypothesis that stimulation of monocytes and macrophages with Colony Stimulating Factor 1 (CSF1) could have a beneficial impact on liver repair after I/R injury.

**Methods:** We investigated the impact of perisurgical treatment with a long-circulating CSF1-Fc conjugate on liver injury and hepatocyte proliferation after 70% ischemia for 60 minutes at 6 h, 48 h and 7 days post reperfusion in male rats. Changes in the circulating and liver tissue monocyte and macrophage subsets in the ischaemic and oxygenated lobes were assessed using quantitative PCR and flow cytometry.

**Results:** CSF1-Fc treatment did not affect the extent of hepatocellular injury post-reperfusion, as indicated by serum transaminases. Liver I/R injury, especially necrotic area, was reduced in CSF1-Fc-treated rats 48 h post-surgery. This was associated with increased accumulation of macrophages in both the oxygenated and ischemic lobes, and localization to necrotic tissue in the ischemic lobe. CSF1-Fc treatment also promoted liver growth, associated with increased parenchymal and non-parenchymal cell proliferation. Flow cytometry and gene expression analysis demonstrated increased monocyte infiltration in ischemic compared to oxygenated lobes. CSF1-Fc increased the abundance of CD43+ non-classical monocytes, consistent with the role of CSF1 signaling in monocyte maturation, and increased CD163 expression on mature macrophages.

**Conclusion:** This study suggests CSF1 stimulation drives monocytes/macrophages towards a pro-regenerative response and perisurgical CSF1 treatment might augment liver regeneration in patients undergoing liver resection.

**Graphical abstract:** 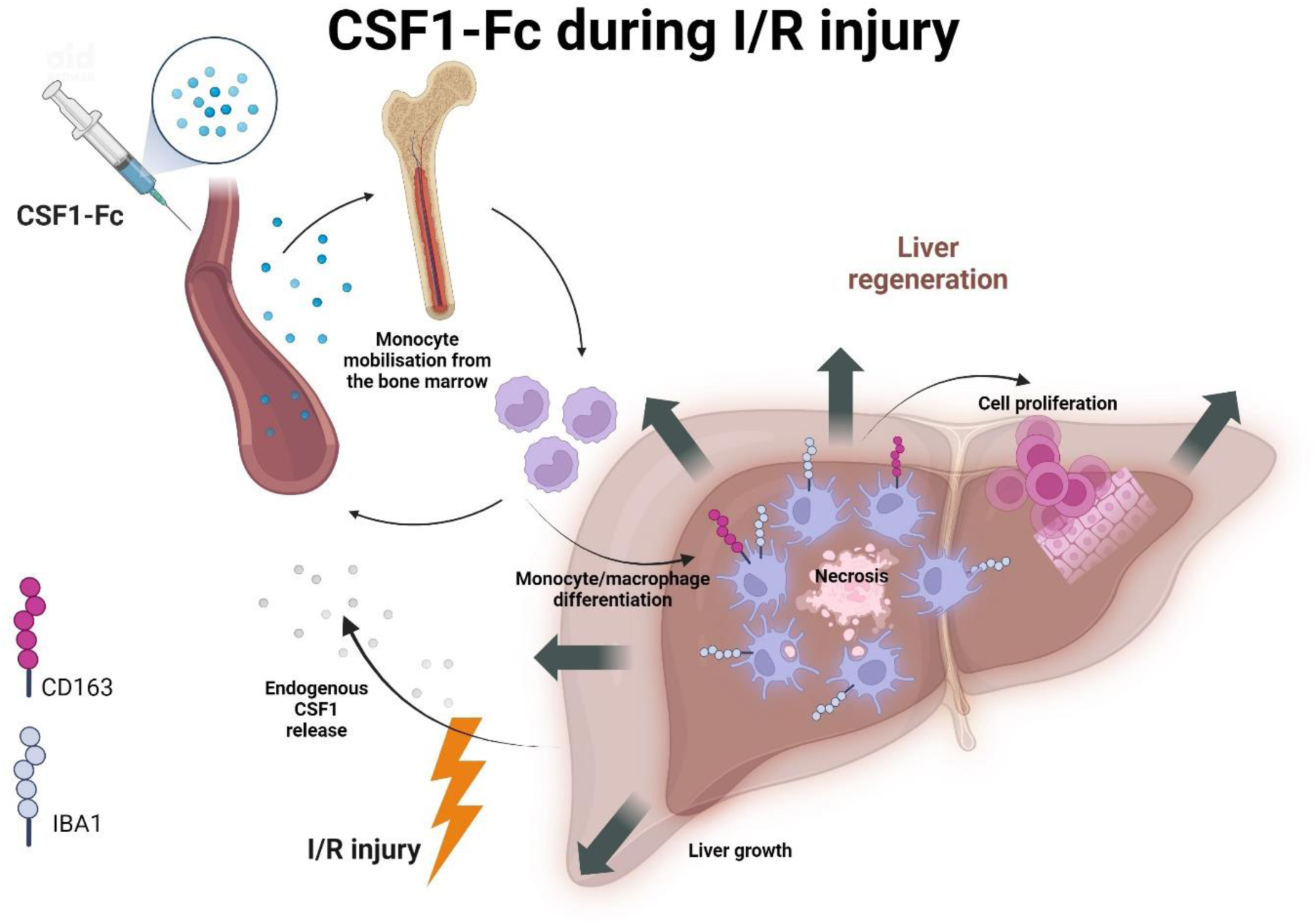

## Introduction

Ischemia and reperfusion (I/R) injury is a well-known complication of surgical vessel clamping, such as the Pringle manoeuvre during hepatectomy to control bleeding. Hepatic I/R injury causes significant morbidity and mortality, especially in organs affected by pre-existing chronic liver diseases and when the interruption to blood flow is prolonged^1^. During hepatic ischaemia, hypoxia and ATP depletion can lead to hepatocyte damage or death, whilst restoration of blood flow triggers oxidative stress and an inflammatory response that can exacerbate tissue damage^2,3^. Although warm and cold ischemia times are strictly controlled in liver transplantation (LTx), I/R injury can lead to severe complications such as graft dysfunction and ischemic-type biliary lesions^4^. Prolongation of ischemic times results in a higher incidence of postreperfusion syndrome and may be associated with poor outcomes and increased risk of HCC recurrence after LTx^5,6^. Recent studies have introduced novel approaches such as the ischemia-free liver transplant (IFLT) technique during which the livers are procured, preserved, and implanted without interruption of normothermic, oxygenated blood supply^7^. While these techniques are not widely performed, have longer organ procurement time and additional costs, hepatic I/R injury remains an ongoing clinical problem.

The pathogenesis of I/R injury involves multiple mechanisms, including oxidative stress caused by reperfusion, and inflammatory cascades mediated by immune cell activation in response to tissue damage. Damage-associated molecular patterns (DAMPs) such as HMGB1, ATP and DNA which are released by ischemia-affected cells, initiating an immune response via pattern recognition receptors (PRRs)^2^. Malfunction of these pathways such as the toll-like receptor 4 (TLR4)-associated activation of JNK and NF-kB, or DNA sensing by cytosolic nucleic acid sensors, compromises the physiological I/R injury response and can impair hepatocyte proliferation and tissue repair^8,9^.

Resident macrophages (Kupffer cells, KC) in the liver contribute to homeostasis and control of immune responses to infection or tissue injury. Macrophages are the earliest responders to I/R injury and orchestrate the subsequent inflammatory response involving monocyte, neutrophil and lymphocyte recruitment and activation. Depending on their activation state, macrophages can both exacerbate inflammation and drive liver disease progression or promote liver repair, e.g. via clearance of damaged cells, matrix remodelling and production of trophic and angiogenic factors and anti-inflammatory cytokines^10,11^. While KC turn over slowly in the steady state and can be maintained by self-renewal, monocyte-derived macrophages (MDM) accumulate in the liver in response to injury-induced chemokines^11^. Resident macrophages and recruited monocytes may play different roles in acute and chronic liver injury, including I/R^12,13^, but both respond dynamically to a rapidly changing microenvironment. We hypothesised that appropriate stimulation of macrophages may be a therapeutic approach to ameliorate I/R injury.

Signalling through the macrophage colony-stimulating factor receptor (CSF1R) drives monocyte differentiation, proliferation, and function^14^. A CSF1-Fc conjugate that has an extended half-life compared to the native protein promoted liver growth in healthy mice, rats and pigs^15-18^, ameliorated acetaminophen-induced acute liver failure^19^, and promoted liver regeneration and fibrosis resolution in mice with toxin-induced fibrosis^20^. Daily CSF1-Fc treatment for 3 days commencing 24 h after hepatic I/R reduced necrotic tissue area 96 h post reperfusion in mice with fibrosis^21^ and CSF1 has also been reported to ameliorate renal I/R^22^. Cell therapy with CSF1-differentiated or alternatively activated macrophages to promote liver regeneration has also been explored^23-25^. On the other hand, depending on dose and time of application relative to injury, CSF1 treatment could exacerbate inflammation and worsen macrophage-mediated pathology in some circumstances^26^. Most of the published studies on CSF1 as a therapy in disease models has been carried out in inbred mice. The rat has many advantages over the mouse, especially for surgical models, and there are significant differences in mononuclear phagocyte biology between the two rodent species, including macrophage expression of CSF1 itself^27^. In this study we tested a regime of perisurgical CSF1-Fc administration as a treatment to promote accumulation of pro-reparative macrophages and ameliorate hepatic I/R injury in rats.

## Results

### Partial liver ischemia reperfusion injury induced endogenous CSF1 24 h post reperfusion

We first conducted a pilot study to determine the time course of liver I/R in Dark Agouti rats. Groups of 4-6 week old rats (n=5/group, mixed gender) were subjected to 70% hepatic ischemia for 45 minutes, to model moderate I/R injury, and sacrificed up to 72 h post reperfusion. On histology, the ischemic lobes (IL) showed different stages of ischemic injury 6-72 h post I/R including zones of confluent coagulative hepatocyte necrosis, in periportal, mid, and pericentral areas. These zones of necrosis show peripheral inflammatory cell accumulation, becoming prominent at 48 h post reperfusion (**Figure 1A**). Ischemic damage was macroscopically apparent on the IL of the liver exposed to I/R (**Figure 1B**). The semi-quantitative Suzuki Score, comprising scores for necrosis, vacuolisation and congestion, was used to assess I/R injury. The Suzuki score peaked at 48 h post reperfusion but was highly variable between animals (**Figure 1C**). There was no clear difference between males and females, but the pilot study was not powered to ascertain gender differences. Circulating liver enzymes AST and ALT peaked 6 h post reperfusion, followed by a gradual decline (**Figure 1D**). Additional hepatobiliary function tests (bilirubin, GGT, AP) were mostly within the reference range, excluding a relevant cholestasis (**Figure 1E**). Endogenous circulating CSF1 was within normal range for wildtype Dark Agouti rats 6 h post I/R, (4-8 ng/ml^27^) and transiently increased 24 h post I/R (**Figure 1F**). Despite the elevated CSF1, the peripheral blood monocyte count was unchanged (**Figure 1G**).

**Fig. 1.**
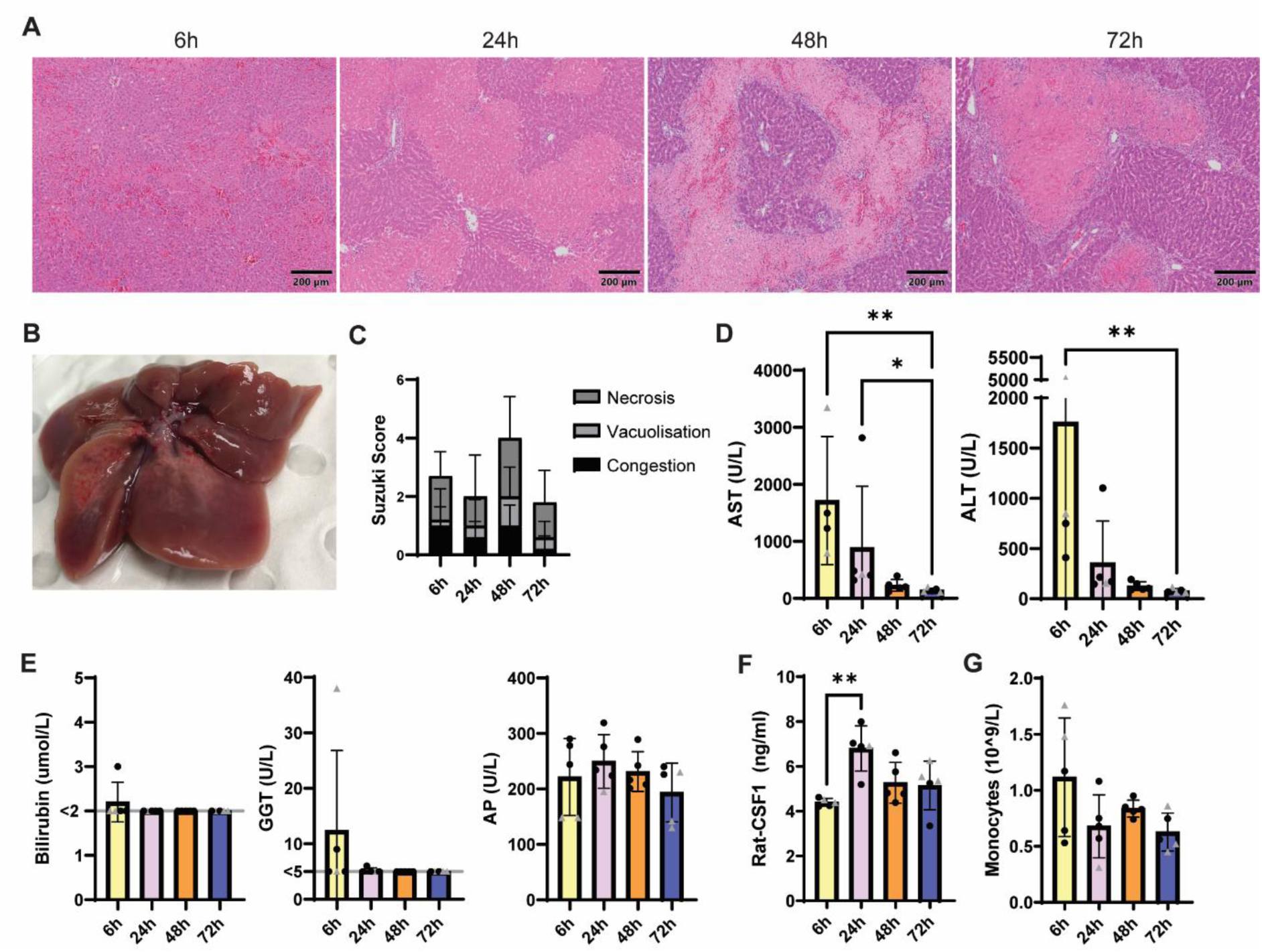
Partial liver ischemia-reperfusion (I/R) injury induced endogenous CSF1 24 h post reperfusion. Groups of 4-6 week old rats (n = 5/group, mixed gender, females indicated by triangle symbol) were subjected to 70% hepatic ischemia for 45 minutes and euthanized at 6, 24, 48 and 72 h post reperfusion. (A) Representative hematoxylin and eosin (H&E) sections of formalin-fixed paraffin-embedded livers showing different stages of ischemic injury at 6-72 h. (B) Macroscopic lesion on the left lateral and median liver lobes at 48 h post I/R. Suzuki Scores (C), serum AST and ALT (D), serum bilirubin, GGT and AP (E). (F) Endogenous serum CSF1 was measured by ELISA. (G) Monocyte count in the peripheral blood was quantified using a haematology analyser. Results were analysed with the Kruskal Wallis test; \**P* < 0.05, \*\*\**P* < 0.001.

### Perisurgical CSF1-Fc treatment increased circulating monocytes and promoted liver growth and repair after ischemic/reperfusion injury

Groups of rats were treated with one daily dose of 1 mg/kg of a human CSF1-mouse Fc conjugate (CSF1-Fc) or saline control on postoperative day -1, 0 and 1 (**Figure 2A**). We initially performed 70% ischaemia for 45 minutes and compared saline and CSF1-Fc-treated rats 48 and 72 h post reperfusion. Liver necrosis had largely resolved by 72 h, especially in CSF1-Fc-treated animals, and we observed high inter-animal variability at 48 h (**Figure S1A**). Others have reported 45 minutes is the approximate threshold of the narrow band of ischemia time that separates negligible from significant I/R injury in mice^28^. Therefore, to reduce inter-animal variability at the peak necrotic timepoint (48 h) we extended the ischaemia time to 60 minutes. Groups of rats were treated CSF1-Fc or saline control as in **Figure 2A** and euthanized 48 h post reperfusion. We previously demonstrated liver regeneration and fibrosis regression after 4 x daily doses of this reagent in mice^20^. In this model CSF1-Fc treatment promoted a small but significant increase in liver/body-weight ratio (**Figure 2B**) but there was no increase in spleen weight (**Figure 2C**). We tested a smaller cohort treated with CSF1-Fc or saline on day -2, -1 and 0 and assessed 6 h post surgery to investigate impacts on the acute response to I/R (**Figure S1B**). In this cohort we did not observe significant changes in liver or spleen/body weight ratio (**Figure S1C-D**). Haematological analysis in both cohorts showed an increase in peripheral blood monocyte count in response to CSF1-Fc injection at 48 h and a trend to increase at 6 h post surgery, whereas the total white blood cell count was not altered (**Figure 2D and S1E**). Circulating granulocytes and lymphocytes increased in response to ischaemic injury, but did not differ between treatment groups (**Figure 2D and S1D**). Acute CSF1-Fc treatment in mice transiently reduced platelet count due to the increased activity of the monocyte/macrophage system, and thrombocytopenia was the dose-limiting toxicity in early clinical studies of CSF1^29-31^. At the dose applied here, the platelet count was reduced in CSF1-Fc-treated rats (**Figure 2E and S1E**), but there was no observable increase in perioperative morbidity. To further categorise the cause of the induced thrombocytopenia, we analysed Mean Platelet Volume (MPV), Platelet Distribution Width (PDW) and Plateletcrit (PCT). While MPV is a known marker of bone marrow platelet production and platelet activation, all three makers have been reported to increase in hyper-destructive causes of thrombocytopenia^32^. CSF1-Fc-treated animals consistently showed a reduced PCT and an increase in PDW (**Figure 2E and S1E**), indicating a reduced volume occupied by platelets in the blood and potentially greater variation in the size of circulating platelets. CSF1-Fc treatment had little impact on MPV. This is consistent with evidence in mice that CSF1 promotes a shortened platelet half-life in the circulation that is compensated by increased platelet production^30^.

**Fig. 2.**
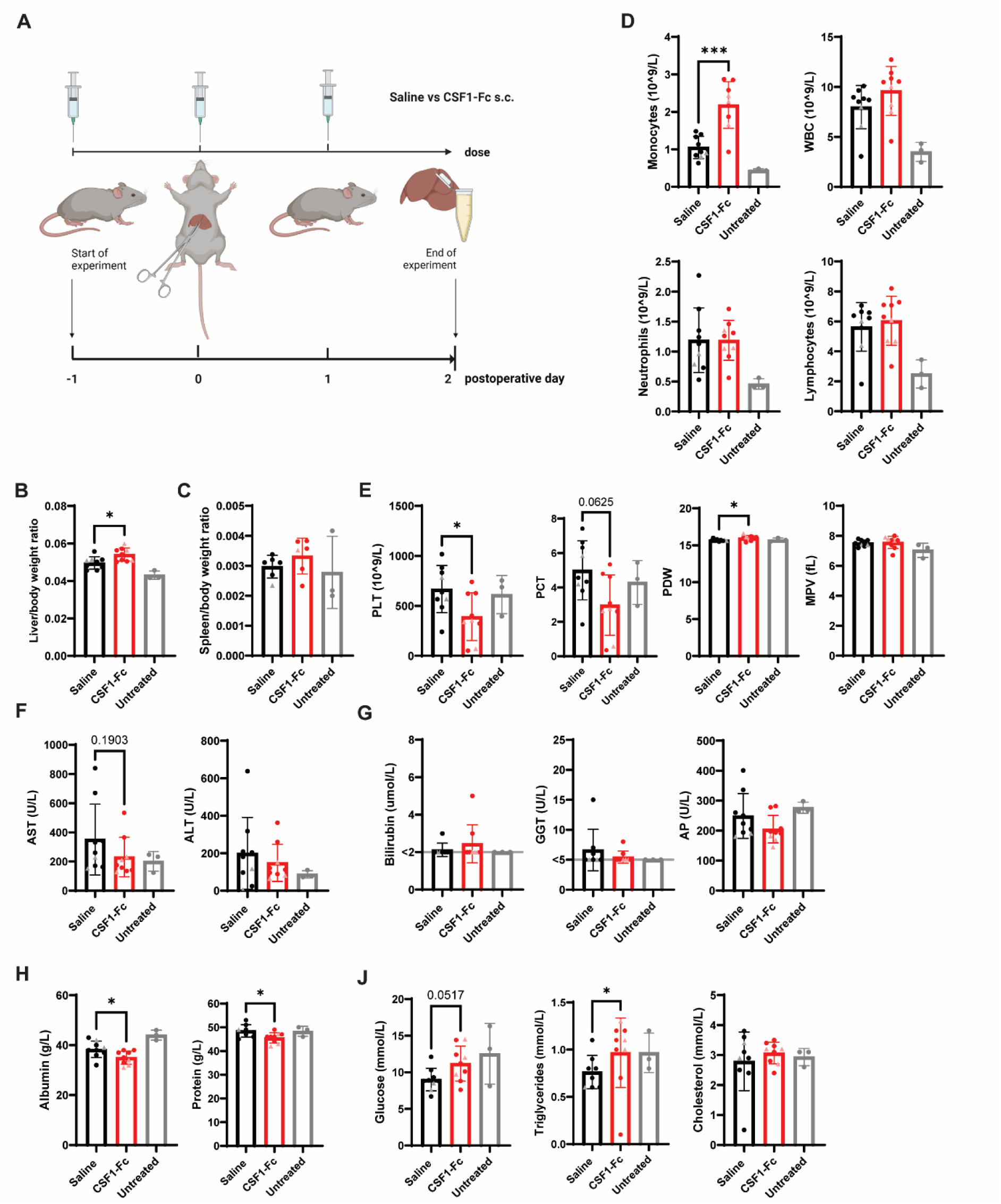
Perisurgical CSF1-Fc treatment increased circulating monocytes and promoted liver growth after I/R injury. (A) Groups of rats (n=9/group except spleen weight n=6/group, females indicated by triangle symbol) were subjected to 60 mins of ischemia, followed by 48 h of reperfusion and treated with 3 x daily doses of 1 mg/kg of a human CSF1-mouse Fc conjugate (CSF1-Fc) or saline control on postoperative day -1, 0 and 1. (B) Liver/body weight ratio. (C) Spleen/body weight ratio. (D) Blood monocyte, white blood cell (WBC), neutrophil and lymphocyte count, (E) platelets (PLT), plateletcrit (PCT), platelet distribution width (PDW) and mean platelet volume (MPV) in the peripheral blood quantified using a haematology analyser. (F) AST and ALT, (G) bilirubin, GGT and AP levels, (H) albumin and total protein and (J) glucose, triglycerides and cholesterol were measured in the serum. Data show mean and standard deviation. Data were analysed using the Mann Whitney test for saline vs CSF1-Fc-treated groups. Data from age-matched Dark Agouti rats (n = 3) that had not been subjected to I/R surgery collected at a different time are shown for reference (grey bars), but not included in statistical analysis.

To address the extent of clinical injury after I/R in control and CSF1-Fc-treated rats, we performed further serum analysis and measured the liver transaminases AST and ALT. Consistent with the pilot data (**Figure 1D**), these injury markers rapidly declined post I/R, but remained modestly elevated above baseline 48 h post surgery. The expansion of blood monocytes in response to CSF1-Fc treatment did not aggravate acute hepatic injury, rather a trend towards reduced levels was apparent, considering the two cohorts together (**Figure 2F and S1F**). Markers of biliary injury (bilirubin, GGT and AP) were mostly within the normal range (**Figure 2G and S1G**), indicating that neither clamping of the bile duct during surgery nor CSF1-Fc treatment caused relevant cholestasis. As previously reported in mice treated with 4 x daily doses of 5 mg/kg human-mouse CSF1-Fc^20^, CSF1-Fc treatment reduced serum albumin and total serum protein (**Figure 2H and S1H**) but had no effect on glucose and cholesterol, but produced a small, significant, increase in triglycerides (TG) at 48h post reperfusion (**Figure 2J and S1J**). Notably, fasting serum cholesterol levels have been reported to decrease during CSF1 infusions^31^, and CSF1-Fc treatment reduced circulating glucose and adiposity in mice^33^. Consistent with the trend to reduced AST and ALT, I/R injury (Suzuki Score) and necrosis area (assessed by image analysis) were significantly reduced in CSF1-Fc-treated animals 48 h post reperfusion (**Figure 3A-B**). Consistent with previous studies^34,35^, I/R did not increase apoptosis in the liver in either the ischemic or oxygenated lobes, as assessed by Caspase 3 activity, nor was there any impact of CSF1-Fc treatment (**Figure 3C**).

**Fig. 3.**
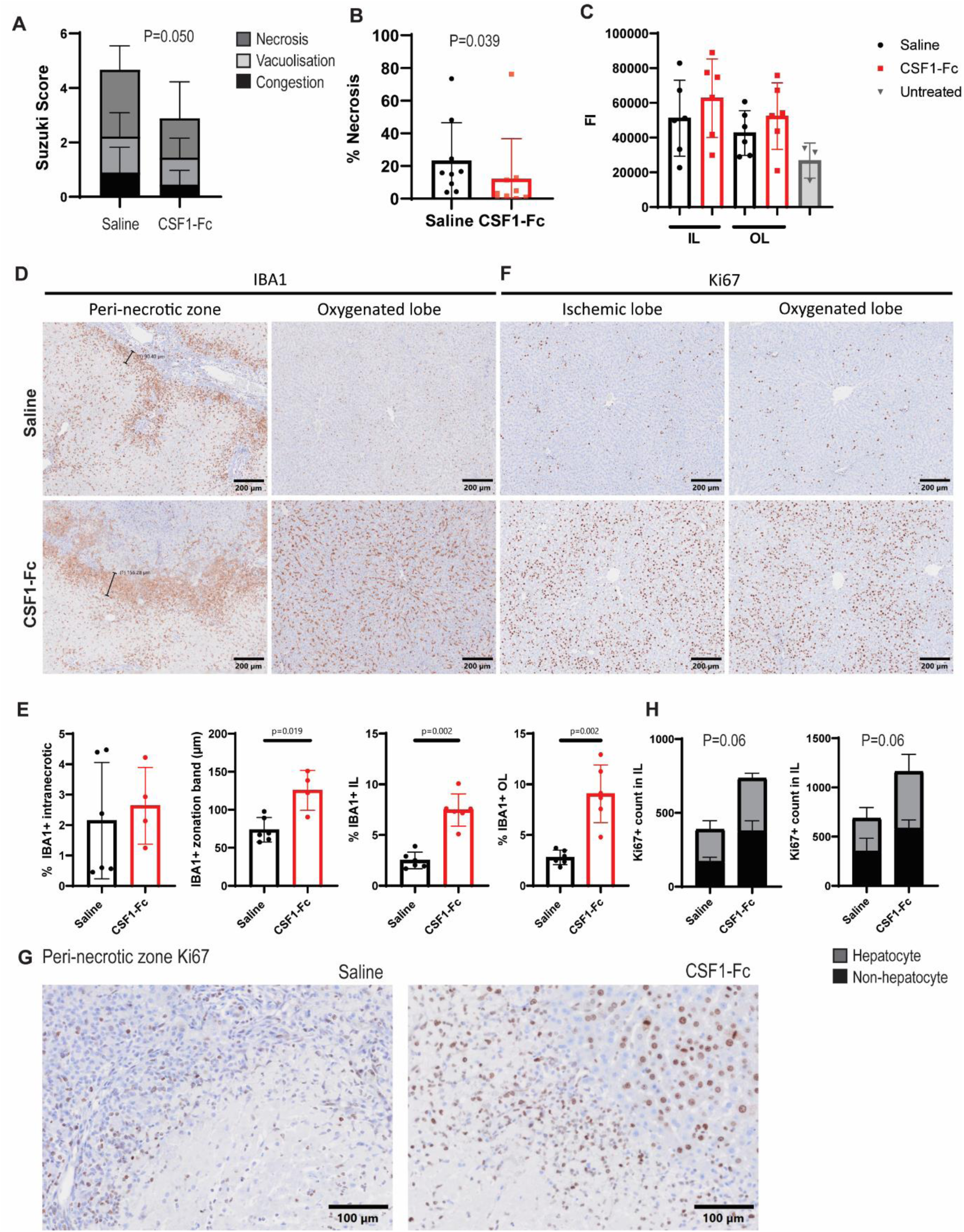
CSF1-Fc treatment reduced the extent of histological necrosis, augmented liver macrophage numbers and promoted cell proliferation. (A) Suzuki Score and (B) necrotic area assessed by image analysis in ischaemic lobes 48 h post reperfusion. (C) Caspase 3 activity in liver lysates, assessed cleavage of a fluorescent substrate (FI = Fluorescent intensity). Liver lysates from age-matched Dark Agouti rats (n = 3) that had not been subjected to I/R surgery collected at a different time were used as untreated controls. Kruskal Wallis test with Dunn’s multiple comparisons test (not significant). (D) Representative immunohistochemistry images of liver IBA1+ cells and (E) quantification of IBA1+ intranecrotic staining area, zonation band thickness and IBA1+ staining area in the ischemic (IL) and oxygenated lobe (OL). The zonation band was not quantified in 3 rats (1 saline, 2 CSF1-Fc) in which no distinct necrotic area was present. (F) Representative immunohistochemistry images of liver Ki67 staining and (G) quantification of Ki67 positive hepatocytes and other cells in the IL and OL. (H) Representative immunohistochemistry images showing Ki67+ cells at a higher magnification in the peri-necrotic zone in the ischemic lobes for saline and CSF1-Fc treatment. Data were analysed using Mann Whitney test, except for (C).

### CSF1-Fc treatment augmented liver macrophage numbers and promoted cell proliferation

To further assess the impact of treatment on I/R pathology sections were stained for the macrophage marker IBA1 and the cell proliferation marker Ki67 (**Figure 3D-G**). A prominent feature of the pathology was the appearance of a peri-necrotic ‘collar’ densely populated with IBA1+ macrophages. Ki67+ cells were present within the necrotic area but were more abundant in the surrounding parenchyma. CSF1-Fc treatment greatly increased the density of IBA1+ cells in both the ischemic and oxygenated lobes and significantly expanded the dense collar of IBA1+ cells surrounding the necrotic zones in the ischemic lobe (**Figure 3D and E**). In line with the increase in liver weight, there was a numerical increase in the numbers of Ki67+ hepatocytes and non-hepatocytes in both the ischaemic and oxygenated lobes in response to CSF1-Fc (p=0.06 for both lobes, **Figures 3F and G**).

To confirm the evidence from IBA1 staining, mRNA was isolated from the livers of control and CSF1-Fc-treated rats and expression of several monocyte/macrophage associated transcripts was measured by qRT-PCR (**Figure 4**). *Adgre1, Cd163, Cd206, Timd4* and *Csf1r* increased in response to CSF1-Fc treatment 48 h after I/R injury, particularly in the OL (**Figure 4A-J**). The type I transmembrane glycoprotein CD163 has been suggested to play a role in resolution of inflammation and regeneration after ischemic injury^36,37^. CD163 is exclusively expressed on macrophages, where it acts as a receptor for haemoglobin:haptoglobin complexes, and is a marker of liver resident macrophages in humans^38^. The effect of CSF1-Fc on *Cd163* mRNA expression appeared greater in the OL compared to the IL (**Figure 4B**). However, increased CD163 expression at the protein level was detected in both ischemic and oxygenated lobes by immunohistochemical staining (**Figure 4N and O**). The induction of CD163 and *Timd4,* a marker of liver resident macrophages in mice and humans that is not expressed on monocyte-derived macrophages, suggests proliferating resident macrophages contribute to the increase in IBA1+ cells (**Figure 4D**). *Mertk*, encoding the efferocytosis receptor MERTK, is another resident macrophage marker in mice and humans, but was not altered by CSF1-Fc treatment (**Figure 4E**). The monocyte chemokine CCL2 is upregulated during development of non-alcoholic steatohepatitis in humans, and in many preclinical models of liver disease^39^, and was highly upregulated by CSF1-Fc in murine liver^20^, but was barely detectable in most lobes in our study (**Figure 4F**).

**Fig. 4.**
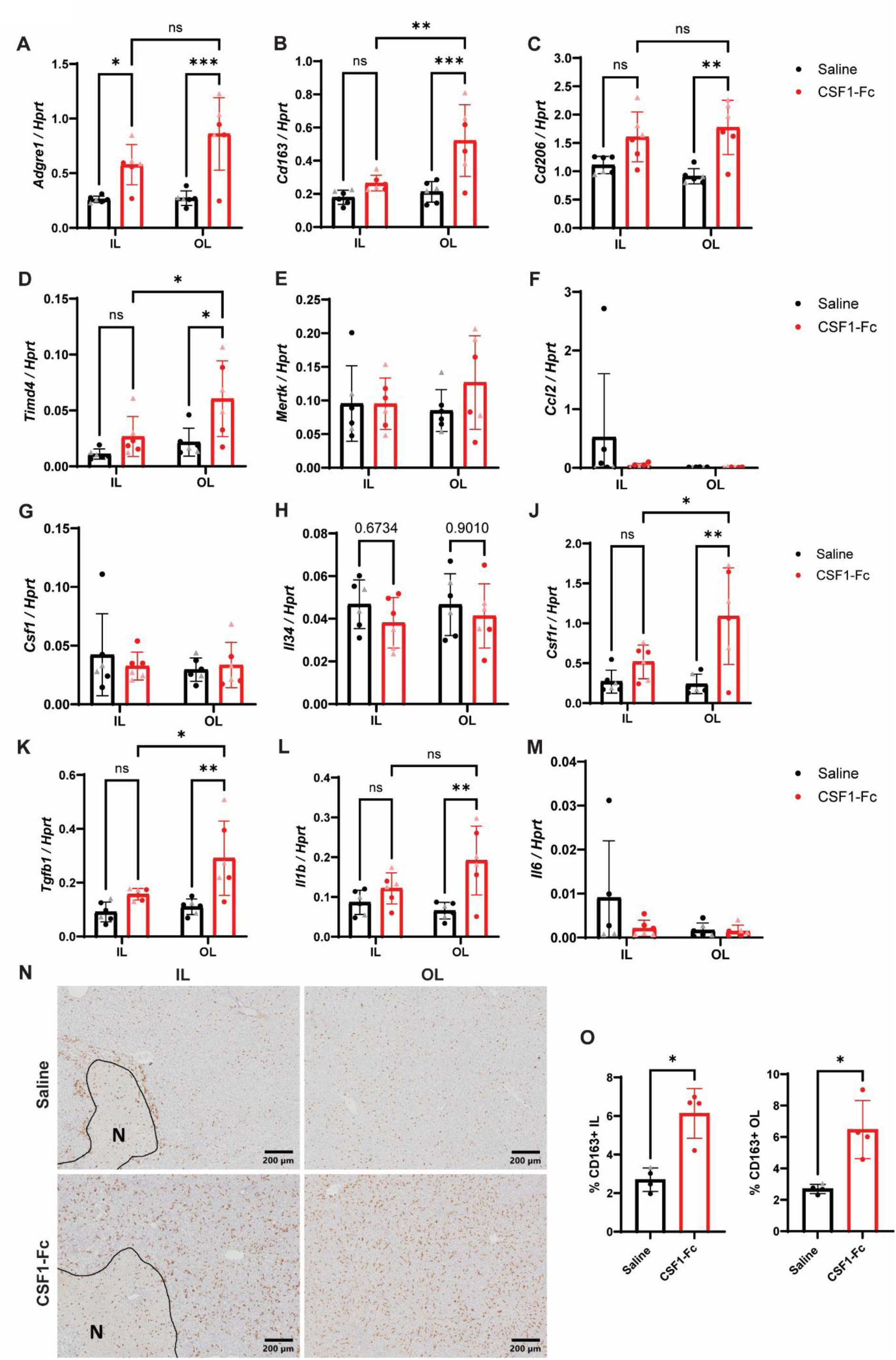
Perisurgical CSF1-Fc treatment increased macrophage and proinflammatory gene expression after liver I/R. (A-M) Whole liver mRNA expression of *Adgre1, Cd163, Cd206, Timd4, Mertk, Ccl2, Csf1, Il34, Csf1r, Tgfb1, Il1b* and *Il6* in the ischemic (IL) and oxygenated lobe (OL) 48 h after I/R. (N) Representative immunohistochemistry images of CD163+ cells and necrosis outlined and indicated with ‘N’). (O) Percentages of CD163+ cells were quantified, n = 4-6 for all groups and results were analysed with ordinary two-way ANOVA with Tukey’s multiple comparisons for A-M and Mann Whitney test for O.

Transcripts encoding both CSF1R ligands, CSF1 and IL34 were detected in rat liver 48 h post reperfusion, but neither was affected by I/R or exogenous CSF1 treatment (**Figure 4G-H**). The early peak in circulating endogenous CSF1 had returned to normal by this timepoint (**Figure 1F**) but this peak may also reflect hepatic clearance. CSF1-Fc treatment upregulated the archetypal profibrotic cytokine transforming growth factor β (*Tgfb1*) mRNA expression in both IL and OL (**Figure 4K**), as occurred in mice^20^. Interleukin-1B (*Il1b*), mRNA expression was also increased in response to CSF1-Fc in the oxygenated lobe, but *Il6*, which was highly inducible in mice^33^, was unaffected (**Figure 4L and M**).

### CSF1-Fc increases non-classical monocytes and CD163+ macrophages in the liver

To further investigate monocyte and macrophage phenotype in liver I/R injury and in response to CSF1-Fc treatment, we analysed cells from disaggregated livers by flow cytometry, utilising the Csf1r-mApple reporter as an additional monocyte/macrophage marker (**Figure 5A and J**). *Csf1r*-mApple rats express a knock-in mApple fluorescent reporter under the control of the *Csf1r* promoter. The reporter is expressed in monocyte/macrophage lineage cells, as well as B cells and neutrophils^16^. The yield of cells from ischemic lobes was significantly higher than from oxygenated lobes, reflecting injury induced inflammation, and CSF1-Fc increased yield in the oxygenated lobe (**Figure 5B**). CD172a (SIRPa) was used to gate myeloid cells (the majority also expressed CD11b/c, not shown), which were further subdivided into CD172Hi monocytes/macrophages and CD172Intermediate neutrophils (which expressed intermediate mApple and HIS48^16^ (not shown)) (**Figure 5A**). CD172aHi monocytes/macrophages followed a similar trend to total cell yield, being the majority of cells recovered, and the CD172a-intermediate neutrophil population increased in the ischemic lobes, regardless of treatment, similar to circulating granulocytes (**Figure 5C and D**). CD43 and HIS48 define monocyte and tissue macrophage subsets in rats^16^. Classical monocytes (the equivalent of Ly6CHi in mouse or CD14Hi in humans) express high levels of HIS48, whereas non-classical monocytes (equivalent of Ly6Clow in mouse or CD16Hi in humans) express high CD43 (**Figure 5A**). CD43Low/HIS48Low cells were gated as macrophages; they expressed the mature macrophage marker CD4, in common with CD43Hi monocytes, and a subset expressed the liver macrophage marker CD163, which was not present on either monocyte population (**Figure 5A**). HIS48Hi classical monocytes increased in the ischemic lobes, especially in saline-treated rats (**Figure 5E**). Ischemia and CSF1-Fc treatment significantly increased the number of CD43Hi monocytes (**Figure 5F**), which may reflect their specific recruitment from the blood (since CD43Hi is the predominant circulating population in rats) or maturation from classical monocytes within the liver. CD43Low/HIS48Low macrophage numbers were non-significantly increased in both ischemic and oxygenated lobes in response to CSF1-Fc treatment (**Figure 5G**), and macrophage CD163 expression also increased (**Figure 5H**), in line with IHC.

**Fig. 5.**
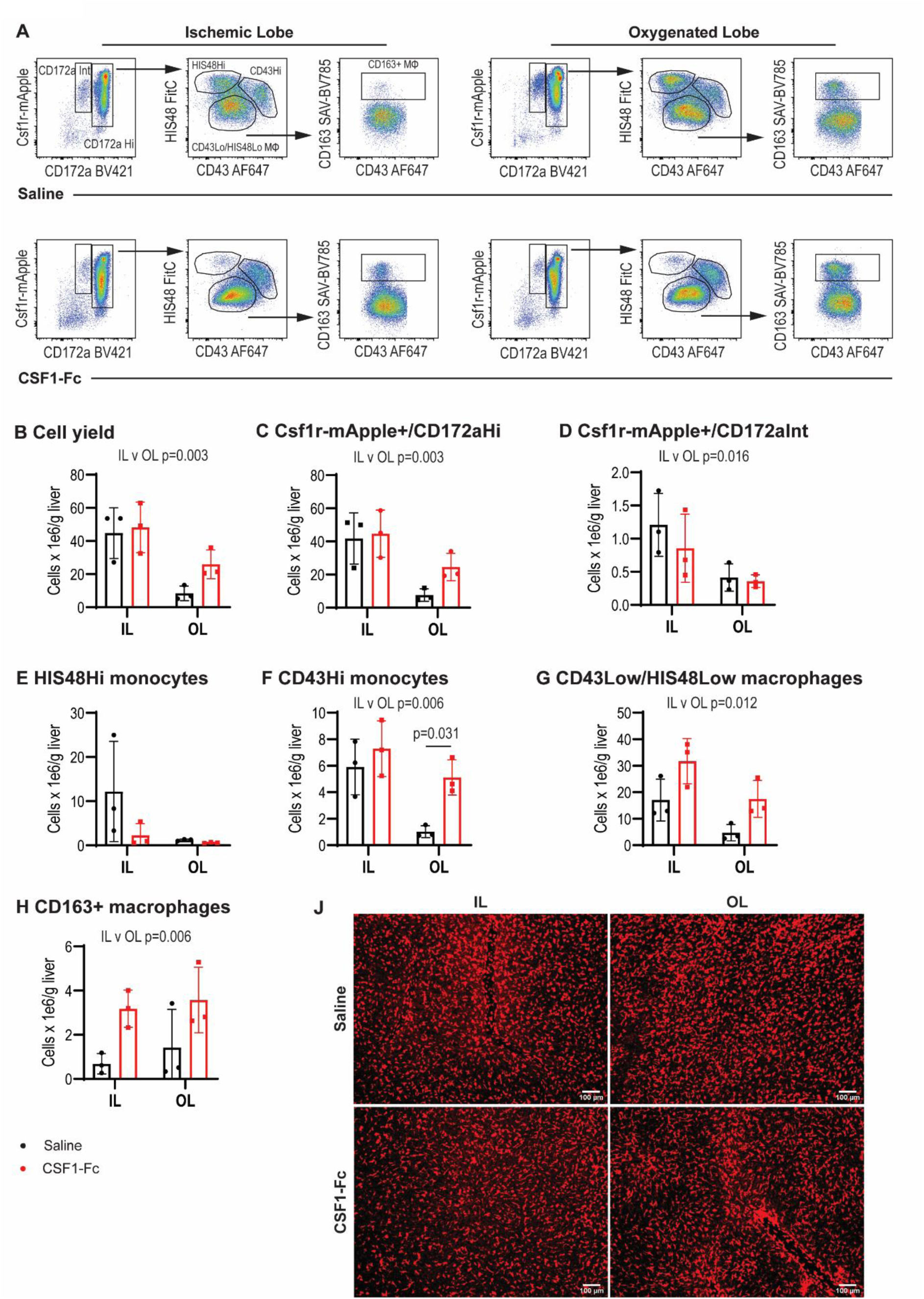
CSF1-Fc mobilises non-classical monocytes into the liver and increases CD163+ macrophages. Non-parenchymal cells were isolated from disaggregated livers harvested from Csf1r-mApple rats 48 h post I/R. (A) Csf1r-mApple expression and CD172a (SIRPa) were used to gate myeloid cells, and further subdivided into CH172Hi (monocytes and macrophages) and CD172Intermediate (neutrophils). CD43Low/HIS48Low cells were gated as macrophages, HIS48Hi as classical monocytes and CD43Hi as non-classical monocytes. The liver macrophage marker CD163 was expressed on a subset of CD43Low/HIS48Low cells and the impact of CSF1 treatment on cell yield and the different subpopulations (B-H) in the ischemic (IL) and oxygenated lobes (OL) were quantified. (J) Representative whole-mount imaging of fresh unfixed tissues from *Csf1r*-mApple transgenic rats 48 h post I/R using a spinning disc confocal microscope, n = 3 for all groups and results were analysed with ordinary two-way ANOVA with Tukey’s multiple comparisons for B-H.

### CSF1-Fc treatment accelerates AST decline after I/R injury

Based on the potential of monocyte-derived macrophages to promote tissue repair^12^, high hepatocyte and non-parenchymal cell proliferative activity, and lack of evidence of adverse impacts, perisurgical CSF1-Fc treatment has the potential to improve long-term recovery from I/R. Therefore, we investigated the outcome of the healing and remodelling phase on postoperative day 7, with CSF1-Fc treatment on days -1, 0, 1 and 2. Due to the variable injury observed, in this experiment we included blood sampling on alternate days post surgery to facilitate analysis of recovery relative to the initial injury in each animal (**Figure 6A**). CSF1-Fc treatment had no impact on the speed of recovery of initial body weight loss (**Figure 6B**). Absolute liver and spleen weight, liver and spleen/body weight ratio and blood monocytes and platelet parameters were at control levels by day 7 (**Figure 6C-G**) consistent with rapid reversal seen in mice^20^. To analyse the resolution of injury we measured AST in the serum obtained from blood draws on postoperative day (POD) 1, 3 and 5. The postoperative decline of AST between POD 1 and 3 (**Figure 6H**) was significantly accelerated by CSF1-Fc treatment, suggesting acute CSF1-Fc treatment promotes tissue repair at least up to 72 h post reperfusion. I/R-induced necrosis was resolved in all animals by day 7, with the exception of 1 saline-treated rat (**Figure 6I**).

**Fig. 6.**
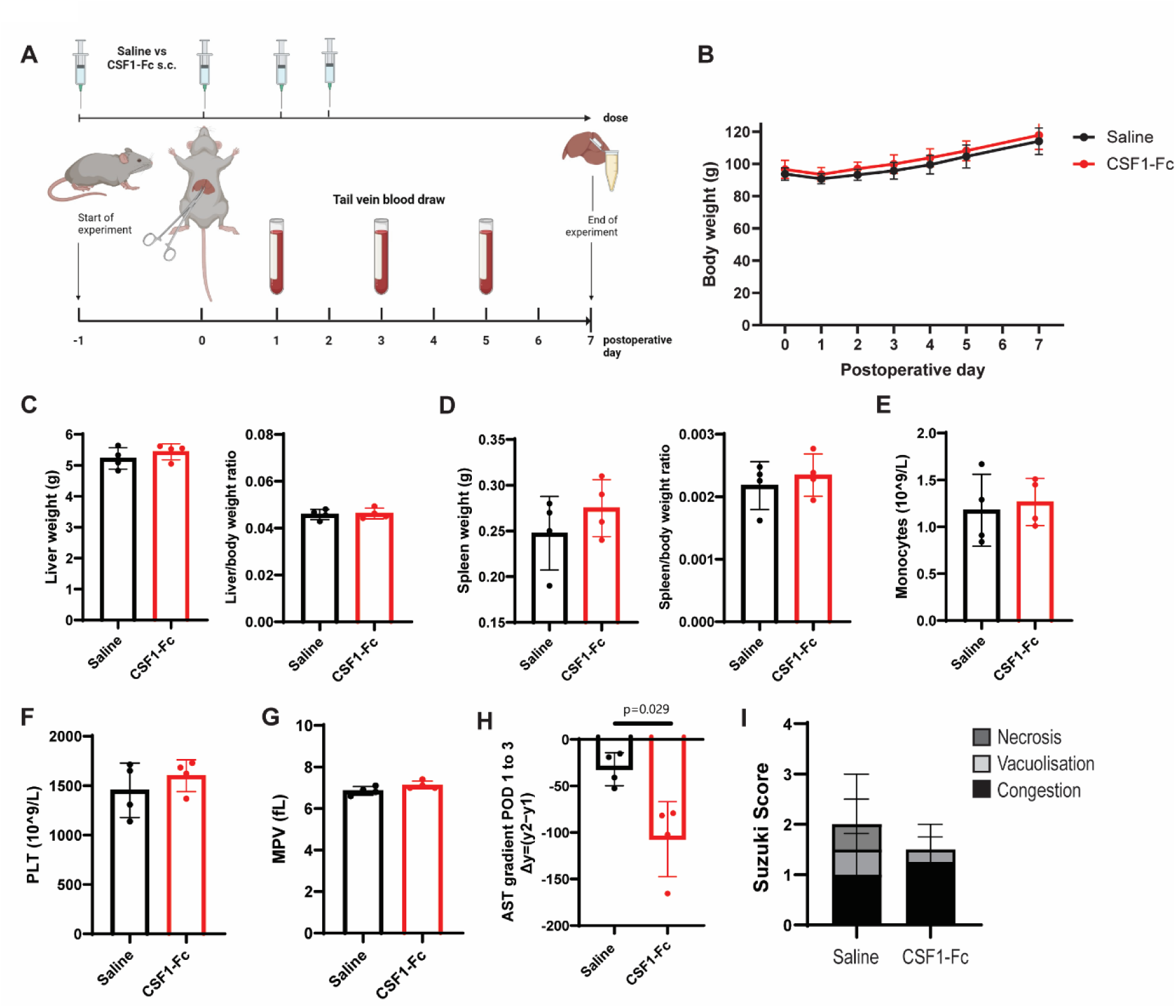
CSF1-Fc treatment accelerates AST decline after I/R injury. (A) Groups of male rats were subjected to 60 mins of ischemia, followed by 7 days of reperfusion and treated with 4 x daily doses of 1 mg/kg of CSF1-Fc or saline control at postoperative day -1, 0, 1 and 2. (B) Postoperative body weight, (C) total liver weight, liver/body weight ratio and (D) total spleen and spleen/body weight ratio were measured. (E) Monocyte count, (F) platelets (PLT) and mean platelet volume (MPV) in the peripheral blood were quantified. (H) AST gradient between postoperative day (POD) 1 to 3. (I) Suzuki scores on postoperative day 7. n = 4 for all groups, results were analysed using the Mann Whitney test.

## Discussion

Monocytes and macrophages have pleiotropic roles in liver injury and repair. Strategies to promote pro-reparative macrophages have significant therapeutic potential in acute and chronic liver injury. The liver-trophic effect of the homeostatic macrophage growth factor CSF1 is well established^15-17,19^, and the transient upregulation of endogenous CSF1 post I/R observed here supports a role in tissue repair. However, CSF1 also has the potential to exacerbate inflammation driven by inflammatory monocytes, depending on the nature of injury and timing of administration^20^. In this study we investigated the impact of perisurgical CSF1-Fc administration on acute I/R injury and tissue repair. Our results indicate that expansion of monocyte and tissue macrophage populations does not exacerbate I/R pathology and that CSF1-Fc treatment has the potential to accelerate the resolution of I/R injury, including tissue necrosis.

The model we used (70% hepatic ischaemia for 45 or 60 minutes) provoked mainly moderate degrees of necrosis, but variability between animals was significant. The narrow time-threshold separating mild from severe injury in rodent models and the importance of empirically establishing an I/R model due to significant inter-laboratory differences have previously been highlighted^28,40,41^. Although prolonged ischemia and severe (>70%) necrosis serve as a more reproducible model, this level of injury is rarely seen in the clinic. Severe necrosis evoked a dominant neutrophil response, whereas mild ischemia triggered a monocyte-driven response, which was more similar to clinical I/R injury^40^. In our rat model, we observed mild-moderate injury that generally resolved spontaneously and elicited a monocyte/macrophage-dominated response. Modest increases in circulating and hepatic granulocytes were observed, but did not differ between treatment groups.

CSF1-Fc treatment increased circulating monocytes and liver macrophages, including the fraction that surrounded necrotic lesions. We observed an altered monocyte and macrophage phenotype, with a shift to the non-classical CD43+ phenotype and induction of CD163, respectively. These direct effects on monocytes/macrophages resulted in increased liver size and reduced Suzuki Score and necrotic area 48 h post I/R. MDMs were also rapidly recruited to the liver in immune-mediated liver injury in mice, where they encapsulated necrotic areas and facilitated repair by promoting hepatocyte survival and necrotic tissue removal^12^. Although the pathogenic mechanisms differ between the two models, we also observed a prominent macrophage collar, which was significantly expanded by CSF1-Fc treatment. These cells were likely monocyte-derived, as we observed an accumulation of IBA1+ but not CD163+ cells. We also noted a trend towards lower circulating transaminases at the acute phase of I/R (6 h) and an accelerated AST clearance between postoperative day 1 and 3, which may be indicative of a beneficial impact on the extent of hepatocellular injury, or simply reflect enhanced clearance of these enzymes by macrophages^42^. Macrophage depletion with clodronate liposomes had little impact on acute I/R injury but did increase circulating transaminases^43^.

The impact of CSF1-Fc on monocyte/macrophage accumulation and hepatocyte proliferation was not restricted to the ischemic lobe. Both gene expression analysis and flow cytometry indicated similar phenotypic changes regardless of injury, with fewer HIS48+ classical monocytes and more CD43Hi monocytes and CD163+ macrophages in CSF1-Fc-treated livers, consistent with the role of CSF1 in driving monocyte and macrophage maturation^44^. Few studies have compared the ischemic and non-ischaemic lobes in hepatic I/R injury. One study noted similar numbers of necrotic cells in the ischaemic and oxygenated lobes after 30 minutes of 70% ischaemia and up to 6 h reperfusion in rats, although no significant necrosis was present after this mild injury^45^. The transcriptomes of the re-perfused lobes were strikingly similar to their non-ischaemic counterparts, which was interpreted as reflecting responses to hemodynamic changes or circulating factors^45^. I/R injury also impaired liver macrophage bacterial uptake and killing capacity to a similar extent in ischaemic and non-ischaemic lobes in mice^46^. Enhanced hepatic growth and macrophage-mediated clearance activity^19^ in unaffected lobes has the potential to mitigate impacts of loss of function in the ischemic lobe and the risk of post-operative infections. Notably, CSF1-Fc treatment did not alter *Mertk*, a marker of liver resident macrophages in humans and mice, consistent with a previous report that this gene is not CSF1-dependent in rats^47^. As observed in mouse liver^20^ CSF1-Fc induced the canonical profibrogenic cytokine *Tgfb1*, which was reported to aggravate hepatic I/R injury^48^. TGFB1 is also a feedback regulator of macrophage differentiation and is produced during spontaneously resolving inflammation in experimental animals, and specifically following the uptake of apoptotic cells by macrophages and may thus also contribute to dampening proinflammatory responses and injury resolution^49^.

Liver I/R injury, as with other liver injuries, may be more severe in male than female patients, and similar findings have been reported in rats^50,51^. The use of mixed gender animal cohorts is a limitation of our study. However, we predominantly used male rats and gender ratios were balanced between treatment groups. We have also previously established that the CSF1-Fc-induced liver regenerative response does not differ between males and females^20^. A larger sample size would be required to confirm previous reports of gender differences in I/R injury.

In summary, our findings suggest supplementary CSF1 therapy during liver resection might facilitate liver regeneration by inducing a pro-reparative macrophage phenotype in ischaemic tissue and augmenting the function of non-injured liver function. These findings provide the basis for future clinical investigations of CSF1-Fc which could lead to the development of new strategies to mitigate I/R injury.

## Material and methods

### Ethics statement

Rats were bred and maintained in specific pathogen-free facilities at The University of Queensland under protocols approved by The University of Queensland Animal Ethics Unit (Approval MRI-UQ/2021/AE000958).

### Animal studies

Csf1r-mApple transgenic reporter rats express mApple fluorescent protein under the control of Csf1r promoter and enhancer elements^16^. They have been back-crossed from the original Sprague-Dawley background to the Dark-Agouti (DA) background, used in this study, for 6 generations. 4-6 week old rats were randomly assigned to timepoint, saline-control and CSF1-Fc treatment groups. Rats in treatment cohorts received 3-4 subcutaneous (s.c.) doses of saline or a human CSF1-mouse Fc conjugate (Novartis, Switzerland) at 1 mg/kg based on previous experiments^16,20^. Partial hepatic I/R surgery was performed by a trained surgeon (SS): Isoflurane anaesthesia was induced and maintained throughout the surgery. After routine pre-surgical preparations (application of eye lubricant, hair clipping) rats were placed supine on a heating pad with anaesthesia maintained via nose cone and with rectal body temperature monitoring. The animal was covered with a sterile plastic drape and a midline laparotomy was performed. The liver lobes were mobilised with sterile cotton tips to expose the portal vein, hepatic artery and bile ducts going to the median and left lobes and clamped by placement of a De Bakey-Hess Bulldog Clamp (BBraun, BH030R). Successful clamping was confirmed by decolouration of the ischemic lobes (IL) vs oxygenated lobes (OL). During ischemia the exposed liver was covered with a sterile gauze pad soaked in warm saline and the gauze was replaced every 15 minutes. The animal was continuously monitored during ischemia, followed by removal of the clamp after 45-60 minutes to initiate reperfusion. Reperfusion was visually verified by restoration of colour to the ischemic lobes (ILs). The incision was closed with a Vicryl 4-0 and Prolene 3-0 (Ethicon) suture and additional wound clips (Clay Adams Autoclip Wound Clip, 427631, BX/1000) for the skin. Postoperative analgesia (0.05 mg/kg Buprenorphine s.c.) was administered twice daily. Mortality rates after surgery were as follows: 3 rats did not survive surgery in the pilot study (respiratory failure, intraoperative bleeding and an unknown cause, respectively), and 6 rats in the treatment cohort died (5 from respiratory failure during surgery and 1 due to ruptured sutures). For the 7 day recovery study, tail vein blood draws were performed under short anaesthesia by isoflurane inhalation on postoperative days 1, 3 and 5. Animals were euthanized by CO2 inhalation. Body weights were recorded, and unfasted blood was collected by cardiac puncture for haematology analysis (Mindray BC-5000) and serum separation. Livers and spleens were weighed and collected for later analysis.

### Histology and confocal microscopy

Livers were fixed in 4% paraformaldehyde for 24 h and paraffin-embedded. 4 μm sections were deparaffinized and rehydrated in a descending ethanol series. For morphological analysis sections were stained with Hematoxylin (Sigma-Aldrich) and Eosin (Eosin Y, Sigma-Aldrich). For immunostaining, epitope retrieval was performed in 10 mM Sodium Citrate Buffer (pH 6.0) in a Biocare Medical Decloaker at 95°C for 20 minutes, followed by staining for Ki67 (Abcam, ab16667, 1:100), IBA1 (Novachem, 019-19741, 1:1000), CD163 (Abcam, ab182422, ERP19518, 1:500). Secondary detection was with DAKO Envision anti-rabbit HRP detection reagents (Agilent Technologies Australia). Sections were counterstained with hematoxylin (Sigma-Aldrich), dehydrated in an ascending ethanol series, clarified with xylene, and mounted with DPX mountant (Sigma-Aldrich). Image quantification was performed from whole-slide digital images (VS120 scanner, Olympus) using ImageJ or Visiopharm software. The semi-quantitative Suzuki Score for I/R injury was determined from haematoxylin and eosin stained sections by a liver pathologist (GM) blinded to treatment groups: Congestion and vacuolisation were scored 0 – 4 as none, minimal, mild, moderate or severe. Necrosis was scored 0 – 4 as none, apoptosis, <30%, 30-60% or > 60%^52^. Necrosis area was also assessed using image analysis of whole slide digital images of H&E-stained sections using Olympus VS200 Desktop 3.3 (CellSens) after training a neural network (VS20S-DNN) to detect necrotic area. The zonation band width analysis of the IBA1+ ‘collar’ was measured in OlyVIA (Olympus) at 5 different sites around the necrotic area for each animal that had developed necrosis and the average was used for statistical analysis. Ki67+ hepatocytes (based on nuclear morphology) and non-hepatocytes were counted using ImageJ in 5 x 20x fields per sample and the average was used for statistical analysis. Direct whole-mount imaging of livers detecting the Csf1r-mApple signal was acquired using the 561-diodelaser on the Laser Scanning Confocal Microscope Olympus FV3000 main combiner (FV31-MCOMB).

### Quantitative measurement of serum CSF1

Serum was separated from whole blood by centrifugation at 5000 g, 4°C for 10 minutes and stored at -80°C. CSF1 was measured using the single-wash 90-minute sandwich Rat M-CSF ELISA Kit (Abcam, ab253214) according to the manufacturer’s instructions.

### Serum biochemical analysis

Aspartate Aminotransferase (AST), Alanine Aminotransferase (ALT), Bilirubin, Gamma Glutamyl Transferase (GGT) and Alkaline Phosphatase (ALP) were measured by the QML Pathology Vetnostics Laboratory (Murarrie QLD, Australia). For small volume serum samples obtained from longitudinal tail vein blood draws in the 7 day recovery we used an AST Colorimetric Activity Assay Kit (Cayman, 701640), as the sample was insufficient for routine chemical pathology analysis. The absorbance was detected using a PHERAstar FSX plate reader (BMG Labtech).

### Caspase 3 activity assay

Liver tissue lysates (∼ 10 mg) were generated in a lysis buffer (50 mM HEPES, pH 7.5, 0.1% CHAPS, 2 mM DTT, 0.1% Nonidet P-40, 1mM EDTA and 1 mM PMFS 1:1000) using 1.4 mm Precellys Zirconium Oxide Beads (Bertin Technologies) in a FastPrep-24 5G Homogenizer (MP Biomedicals) for 40 seconds at 4 m/s. Protein lysates were incubated on ice for 15 mins before protein concentration was measured by BCA Assay. Caspase Assay Buffer (100 mM HEPES, pH 7.2, 10% sucrose, 0.1% CHAPS, 1 mM Na-EDTA and 2mM DTT) was added to tissue lysates at equal protein concentrations. The fluorogenic caspase 3 substrate Ac-DEVD-AMC (Cayman, 14986) (10 mM) was dissolved in DMSO and added to initiate the reaction, followed by an incubation for 1 h at 37 °C. Signal intensity was read at excitation 380 nm and emission at 460 nm using a PHERAstar FSX plate reader (BMG Labtech).

### Flow cytometry

Liver non-parenchymal cells (NPCs) were isolated by tissue disaggregation of finely chopped liver samples (∼1-2 g) in 10 ml digestion solution containing 1 mg/ml Collagenase IV (Gibco, 17104-019), 0.4 mg/ml Dispase II (Gibco, 17105-041) and 20 μg/ml DNAse1 (Roche) in 1X HBSS (Gibco, 14065-056) and incubating at 37°C for 45 min on a rocking platform before mashing through a 70 μm filter (Falcon). Hepatocytes were extracted by two runs of a slow spin (50 g) and the supernatant enriched for NPCs was pelleted and red blood cell lysis was performed in ACK lysis buffer for 2 minutes (150 mM NH4Cl, 10 mMKHCO3, 0.1 mM EDTA, pH 7.4). NPCs were centrifuged and washed twice in PBS, and the pellet was resuspended in FC buffer (PBS/2% FBS) for staining. Mononuclear cells were counted using a Mindray analyser. Cells were stained for 30 min on ice in FC buffer containing unlabelled CD32 (BD Biosciences) to block CD32 Fc receptor binding, HIS48-FITC, CD11B/C-BV510, CD45-PE/Cy7, CD172A-BV421 (BD Biosciences), CD43-AF647, CD4-APC/Cy7, (BioLegend), and CD163-Biotin (Bio-Rad). After primary antibody staining cells were washed and stained with streptavidin SAV-BV785 (BioLegend) for 30 min on ice in FC buffer. Cells were washed twice and resuspended in FC buffer containing 7AAD (LifeTechnologies) for acquisition on a Cytoflex (Beckman Coulter LifeSciences). Single-color controls were used for compensation and unstained cells were used to confirm gating strategies. FC data were analysed using FlowJo 10 (Tree Star). Live single cells were identified for phenotypic analysis by excluding doublets (FSC-A>FSC-H), 7AAD+dead cells, and debris. Cell counts were calculated by multiplying the frequency of the cell type of interest by the total mononuclear cell yield/g of disaggregated tissue.

### qPCR

Liver samples were collected in TRIzol (Invitrogen, 15596026), homogenized with 1.4 mm Precellys Zirconium Oxide Beads (Bertin Technologies) for 30 seconds at 4 m/s in a FastPrep-24 5G Homogenizer (MP Biomedicals), followed by stepwise RNA extraction according to the user instructions. cDNA synthesis was performed after recombinant DNase I treatment (Roche) using the SensiFAST cDNA synthesis mix (Bioline) according to manufacturer’s instructions. RT-PCR was performed using the SYBR Select Master Mix (Thermo Fisher Scientific) on an Applied Biosystems system with default cycling conditions. The following primer pairs were used in this study (Table 1).

**Table 1:**
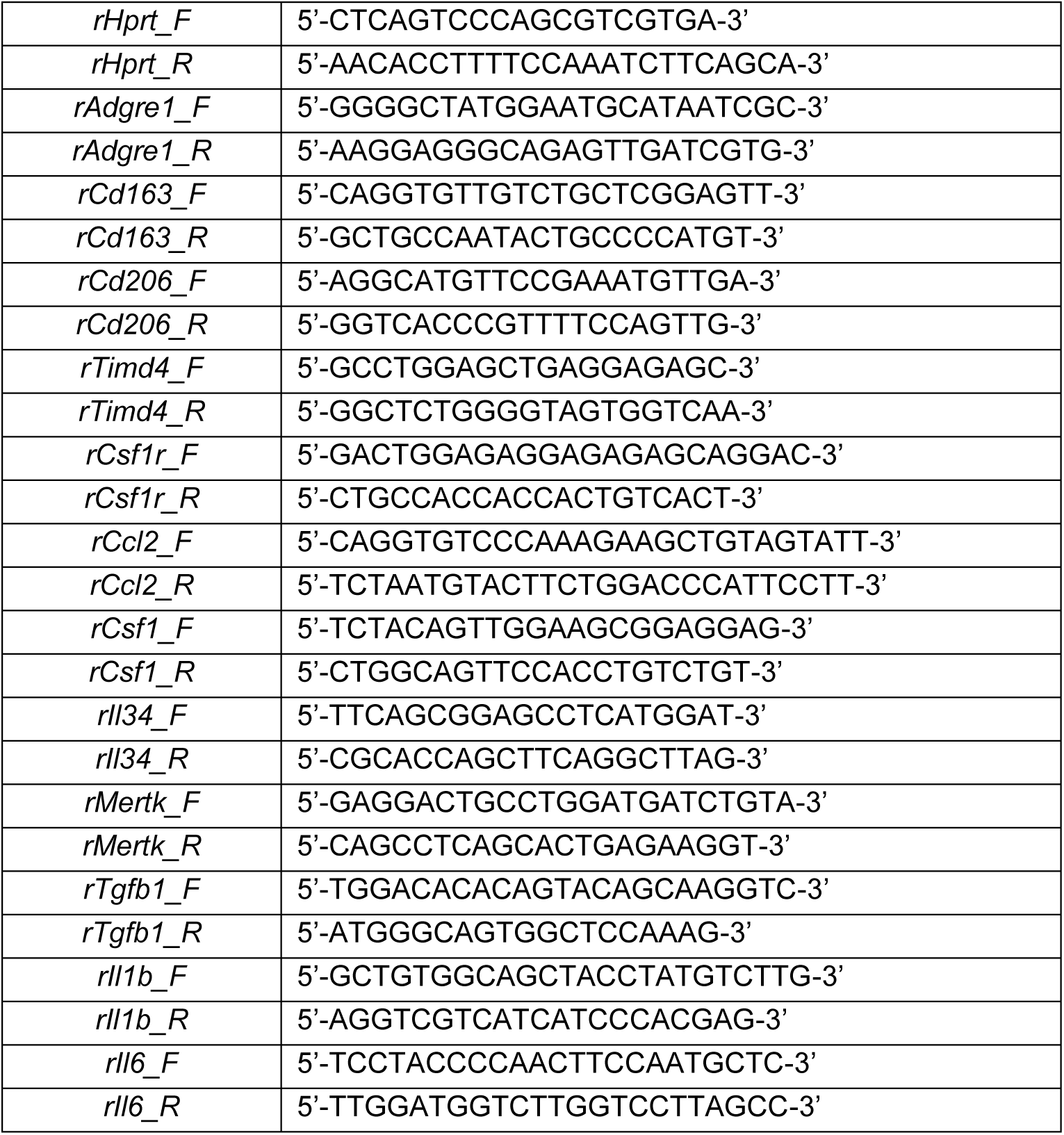

### Statistical Analysis

Analysis of histological and flow cytometry outcome data was performed blinded to treatment group. Data are presented as mean±SD. Statistical tests were performed using GraphPad Prism 8.3.1. Data were analysed using the Mann-Whitney test, Kruskal Wallis test or ordinary 2-way ANOVA, as indicated in Figure legends.

## Acknowledgement

This work was funded by the German Research Foundation (Deutsche Forschungsgemeinschaft) with the Walter Benjamin fellowship project-ID: 490753453 to S.S., the NHMRC Ideas 2012793 to K.M.I. and NHMRC Investigator Grant 2009750 to D.A.H. We appreciate the core laboratory support from the Mater Foundation, the support of The University of Queensland, UQ-PACE Biological Resources Facility, Histology, Microscopy and Flow Cytometry Core Facilities at the Translational Research Institute. We are grateful to Dr Jennifer Borowsky, Pathology Queensland, for helpful discussions. Schematic figures were prepared using Biorender.

## Conflict of interest

The authors declare no competing or financial interests.

## Figures and Legends

**Supp. Fig.1.**
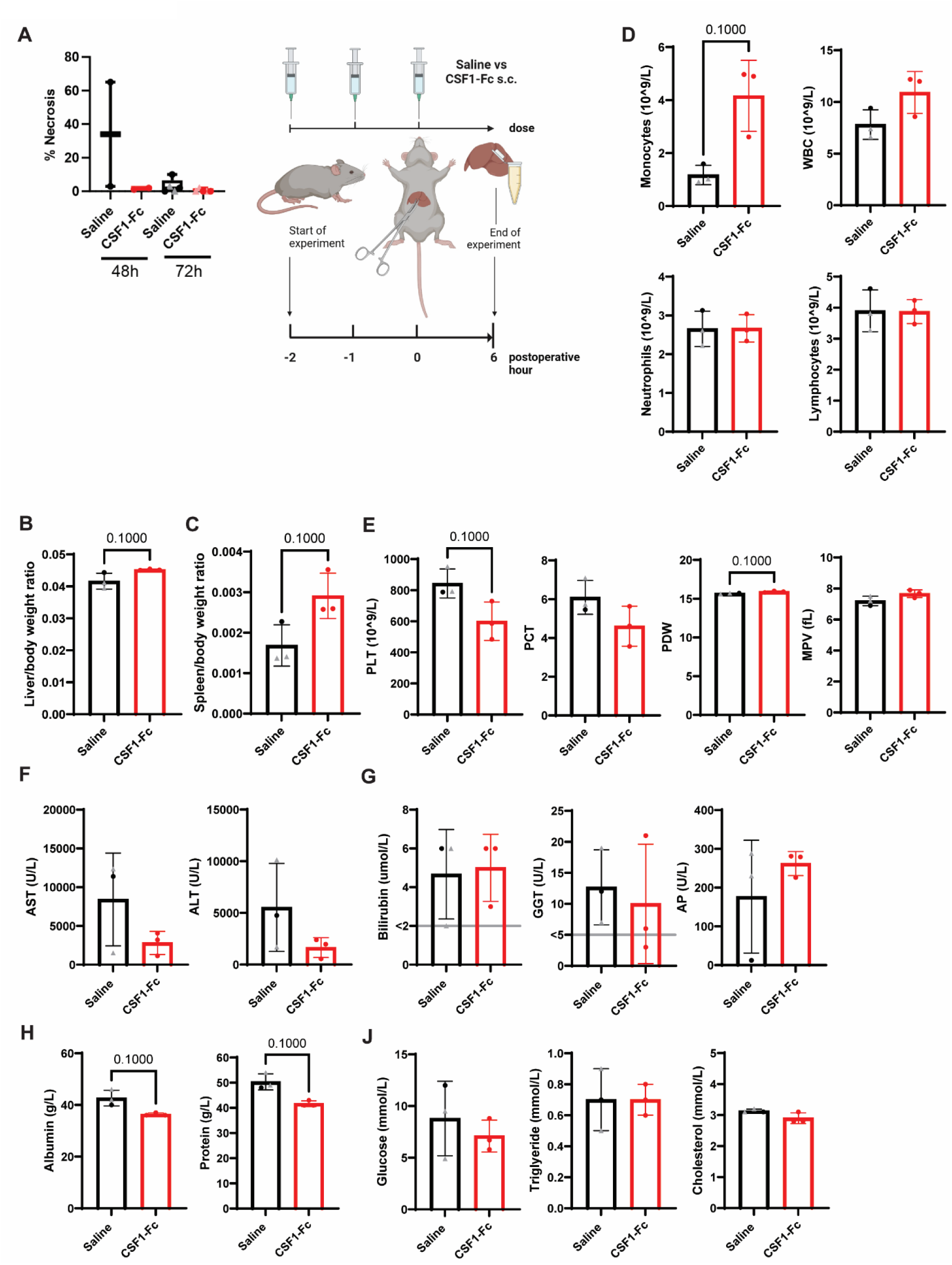
The impact of CSF1-Fc treatment on acute I/R injury. (A) Groups of 2-5 rats (mixed gender, females indicated by triangle symbol) were treated with 3 x daily doses of 1 mg/kg of CSF1-Fc or saline control on postoperative day -1, 0 and 1. On day 0 they were subjected to 45 mins of ischemia, followed by 48 or 72 h of reperfusion. Necrotic area in the ischaemic lobe, box plots with line at median. (B) Groups of male rats were subjected to 60 mins of ischemia, followed by 6 h of reperfusion and treated with 3 x daily doses of 1 mg/kg of CSF1-Fc or saline control at postoperative day -2, -1 and 0 (at the time of surgery). Created with BioRender.com. (B) Total liver weight and liver/body weight ratio and (C) total spleen weight and spleen/body weight ratio were measured. (D) Blood monocyte count, white blood cell (WBC), neutrophil and lymphocyte count, (E) platelets (PLT), plateletcrit (PCT), platelet distribution width (PDW) and mean platelet volume (MPV) in the peripheral blood were quantified using a haematology analyser. (F) AST and ALT, (G) bilirubin, GGT and AP levels, (H) albumin and total protein and (J) glucose, triglycerides and cholesterol were measured in the serum; n = 3 for Saline vs CSF1, and results were analysed using the Mann Whitney test.

## Notes

### Competing Interest Statement

The authors have declared no competing interest.

## References

1. Ishizaki Y, Yoshimoto J, Miwa K, Sugo H, Kawasaki S. Safety of prolonged intermittent pringle maneuver during hepatic resection. Archives of surgery (Chicago, Ill : 1960). 2006;141(7):649-653; discussion 654.

2. Kaltenmeier C, Wang R, Popp B, Geller D, Tohme S, Yazdani HO. Role of Immuno-Inflammatory Signals in Liver Ischemia-Reperfusion Injury. Cells. 2022;11(14).

3. Mao XL, Cai Y, Chen YH, et al. Novel Targets and Therapeutic Strategies to Protect Against Hepatic Ischemia Reperfusion Injury. Front Med (Lausanne). 2021;8:757336.

4. Karp SJ, Johnson S, Evenson A, et al. Minimising cold ischaemic time is essential in cardiac death donor-associated liver transplantation. HPB : the official journal of the International Hepato Pancreato Biliary Association. 2011;13(6):411–416.

5. Peralta C, Jiménez-Castro MB, Gracia-Sancho J. Hepatic ischemia and reperfusion injury: effects on the liver sinusoidal milieu. Journal of hepatology. 2013;59(5):1094–1106.

6. Kornberg A, Witt U, Kornberg J, Friess H, Thrum K. Extended Ischemia Times Promote Risk of HCC Recurrence in Liver Transplant Patients. Digestive diseases and sciences. 2015;60(9):2832–2839.

7. Guo Z, Zhao Q, Jia Z, et al. A randomized-controlled trial of ischemia-free liver transplantation for end-stage liver disease. Journal of hepatology. 2023;79(2):394–402.

8. Schulze S, Stöß C, Lu M, et al. Cytosolic nucleic acid sensors of the innate immune system promote liver regeneration after partial hepatectomy. Scientific reports. 2018;8(1):12271.

9. Tsung A, Hoffman RA, Izuishi K, et al. Hepatic ischemia/reperfusion injury involves functional TLR4 signaling in nonparenchymal cells. Journal of immunology (Baltimore, Md : 1950). 2005;175(11):7661-7668.

10. Duffield JS, Forbes SJ, Constandinou CM, et al. Selective depletion of macrophages reveals distinct, opposing roles during liver injury and repair. The Journal of clinical investigation. 2005;115(1):56–65.

11. Tacke F. Targeting hepatic macrophages to treat liver diseases. J Hepatol. 2017;66(6):1300–1312.

12. Feng D, Xiang X, Guan Y, et al. Monocyte-derived macrophages orchestrate multiple cell-type interactions to repair necrotic liver lesions in disease models. The Journal of clinical investigation. 2023;133(15).

13. Yue S, Zhou H, Wang X, Busuttil RW, Kupiec-Weglinski JW, Zhai Y. Prolonged Ischemia Triggers Necrotic Depletion of Tissue-Resident Macrophages To Facilitate Inflammatory Immune Activation in Liver Ischemia Reperfusion Injury. J Immunol. 2017;198(9):3588–3595.

14. Hume DA, Irvine KM, Pridans C. The Mononuclear Phagocyte System: The Relationship between Monocytes and Macrophages. Trends Immunol. 2019;40(2):98–112.

15. Gow DJ, Sauter KA, Pridans C, et al. Characterisation of a novel Fc conjugate of macrophage colony-stimulating factor. Mol Ther. 2014;22(9):1580–1592.

16. Irvine KM, Caruso M, Cestari MF, et al. Analysis of the impact of CSF-1 administration in adult rats using a novel Csf1r-mApple reporter gene. J Leukoc Biol. 2020;107(2):221–235.

17. Sauter KA, Waddell LA, Lisowski ZM, et al. Macrophage colony-stimulating factor (CSF1) controls monocyte production and maturation and the steady-state size of the liver in pigs. Am J Physiol Gastrointest Liver Physiol. 2016;311(3):G533–547.

18. Pridans C, Sauter KA, Irvine KM, et al. Macrophage colony-stimulating factor increases hepatic macrophage content, liver growth, and lipid accumulation in neonatal rats. Am J Physiol Gastrointest Liver Physiol. 2018;314(3):G388–G398.

19. Stutchfield BM, Antoine DJ, Mackinnon AC, et al. CSF1 Restores Innate Immunity After Liver Injury in Mice and Serum Levels Indicate Outcomes of Patients With Acute Liver Failure. Gastroenterology. 2015;149(7):1896–1909.e1814.

20. Keshvari S, Genz B, Teakle N, et al. Therapeutic potential of macrophage colony-stimulating factor in chronic liver disease. Disease models & mechanisms. 2022;15(4).

21. Konishi T, Schuster RM, Goetzman HS, Caldwell CC, Lentsch AB. Fibrotic liver has prompt recovery after ischemia-reperfusion injury. Am J Physiol Gastrointest Liver Physiol. 2020;318(3):G390–G400.

22. Alikhan MA, Jones CV, Williams TM, et al. Colony-stimulating factor-1 promotes kidney growth and repair via alteration of macrophage responses. The American journal of pathology. 2011;179(3):1243–1256.

23. Dwyer BJ, Macmillan MT, Brennan PN, Forbes SJ. Cell therapy for advanced liver diseases: Repair or rebuild. J Hepatol. 2021;74(1):185–199.

24. Thomas JA, Pope C, Wojtacha D, et al. Macrophage therapy for murine liver fibrosis recruits host effector cells improving fibrosis, regeneration, and function. Hepatology (Baltimore, Md). 2011;53(6):2003–2015.

25. Starkey Lewis P, Campana L, Aleksieva N, et al. Alternatively activated macrophages promote resolution of necrosis following acute liver injury. J Hepatol. 2020;73(2):349–360.

26. Hume DA, MacDonald KP. Therapeutic applications of macrophage colony-stimulating factor-1 (CSF-1) and antagonists of CSF-1 receptor (CSF-1R) signaling. Blood. 2012;119(8):1810–1820.

27. Hume DA, Caruso M, Keshvari S, et al. The Mononuclear Phagocyte System of the Rat. Journal of immunology (Baltimore, Md : 1950). 2021;206(10):2251-2263.

28. van Golen RF, Reiniers MJ, Heger M, Verheij J. Solutions to the discrepancies in the extent of liver damage following ischemia/reperfusion in standard mouse models. Journal of hepatology. 2015;62(4):975–977.

29. Motoyoshi K. Macrophage colony-stimulating factor for cancer therapy. Oncology. 1994;51(2):198–204.

30. Baker GR, Levin J. Transient thrombocytopenia produced by administration of macrophage colony-stimulating factor: investigations of the mechanism. Blood. 1998;91(1):89–99.

31. Jakubowski AA, Bajorin DF, Templeton MA, et al. Phase I study of continuous-infusion recombinant macrophage colony-stimulating factor in patients with metastatic melanoma. Clin Cancer Res. 1996;2(2):295–302.

32. Saran K, Vidya K, Seema K, Prasad A, Prakash J. Study of platelet indices and their role in evaluation of thrombocytopenia. Journal of family medicine and primary care. 2022;11(10):6236–6242.

33. Keshvari S, Masson JJR, Ferrari-Cestari M, et al. Reversible expansion of tissue macrophages in response to macrophage colony-stimulating factor (CSF1) transforms systemic lipid and carbohydrate metabolism. American journal of physiology Endocrinology and metabolism. 2023.

34. Jaeschke H, Lemasters JJ. Apoptosis versus oncotic necrosis in hepatic ischemia/reperfusion injury. Gastroenterology. 2003;125(4):1246–1257.

35. Gujral JS, Bucci TJ, Farhood A, Jaeschke H. Mechanism of cell death during warm hepatic ischemia-reperfusion in rats: apoptosis or necrosis? Hepatology (Baltimore, Md). 2001;33(2):397–405.

36. Akahori H, Karmali V, Polavarapu R, et al. CD163 interacts with TWEAK to regulate tissue regeneration after ischaemic injury. Nature communications. 2015;6:7792.

37. Fabriek BO, Dijkstra CD, van den Berg TK. The macrophage scavenger receptor CD163. Immunobiology. 2005;210(2-4):153–160.

38. Guilliams M, Bonnardel J, Haest B, et al. Spatial proteogenomics reveals distinct and evolutionarily conserved hepatic macrophage niches. Cell. 2022;185(2):379–396 e338.

39. Marra F, Tacke F. Roles for chemokines in liver disease. Gastroenterology. 2014;147(3):577–594.e571.

40. Olthof PB, van Golen RF, Meijer B, et al. Warm ischemia time-dependent variation in liver damage, inflammation, and function in hepatic ischemia/reperfusion injury. Biochimica et biophysica acta Molecular basis of disease. 2017;1863(2):375–385.

41. Karatzas T, Neri AA, Baibaki ME, Dontas IA. Rodent models of hepatic ischemia-reperfusion injury: time and percentage-related pathophysiological mechanisms. The Journal of surgical research. 2014;191(2):399–412.

42. Pervin M, Golbar HM, Bondoc A, Izawa T, Kuwamura M, Yamate J. Immunophenotypical characterization and influence on liver homeostasis of depleting and repopulating hepatic macrophages in rats injected with clodronate. Experimental and toxicologic pathology : official journal of the Gesellschaft fur Toxikologische Pathologie. 2016;68(2-3):113–124.

43. Reiling J, Bridle KR, Schaap FG, et al. The role of macrophages in the development of biliary injury in a lipopolysaccharide-aggravated hepatic ischaemia-reperfusion model. Biochimica et biophysica acta Molecular basis of disease. 2018;1864(4 Pt B):1284-1292.

44. Keshvari S, Caruso M, Teakle N, et al. CSF1R-dependent macrophages control postnatal somatic growth and organ maturation. PLoS genetics. 2021;17(6):e1009605.

45. Zabala V, Boylan JM, Thevenot P, et al. Transcriptional changes during hepatic ischemia-reperfusion in the rat. PloS one. 2019;14(12):e0227038.

46. Klein AS, Zhadkevich M, Wang D, Margolick JB, Winkelstein JA, Bulkley GB. Discriminant quantitation of posttransplant hepatic reticuloendothelial function. The impact of ischemic preservation. Transplantation. 1996;61(8):1156–1161.

47. Pridans C, Irvine KM, Davis GM, Lefevre L, Bush SJ, Hume DA. Transcriptomic Analysis of Rat Macrophages. Frontiers in immunology. 2020;11:594594.

48. Li H, Shen X, Tong Y, et al. Aggravation of hepatic ischemia-reperfusion injury with increased inflammatory cell infiltration is associated with the TGF-β/Smad3 signaling pathway. Molecular medicine reports. 2021;24(2).

49. Sakai M, Troutman TD, Seidman JS, et al. Liver-Derived Signals Sequentially Reprogram Myeloid Enhancers to Initiate and Maintain Kupffer Cell Identity. Immunity. 2019;51(4):655–670.e658.

50. Lu P, Liu F, Wang CY, et al. Gender differences in hepatic ischemic reperfusion injury in rats are associated with endothelial cell nitric oxide synthase-derived nitric oxide. World J Gastroenterol. 2005;11(22):3441–3445.

51. Li Z, Lu S, Qian B, et al. Sex differences in hepatic ischemia‒reperfusion injury: a cross-sectional study. Scientific reports. 2023;13(1):5724.

52. Suzuki S, Toledo-Pereyra LH, Rodriguez FJ, Cejalvo D. Neutrophil infiltration as an important factor in liver ischemia and reperfusion injury. Modulating effects of FK506 and cyclosporine. Transplantation. 1993;55(6):1265-1272.

